# Discovery of a SARS-CoV-2 Broadly-Acting Neutralizing Antibody with Activity against Omicron and Omicron + R346K Variants

**DOI:** 10.1101/2022.01.19.476998

**Authors:** J. Andrew Duty, Thomas Kraus, Heyue Zhou, Yanliang Zhang, Namir Shaabani, Soner Yildiz, Na Du, Alok Singh, Lisa Miorin, Donghui Li, Karen Stegman, Sabrina Ophir, Xia Cao, Kristina Atanasoff, Reyna Lim, Shreyas Kowdle, Juan Manuel Carreño, Laura Rivero-Nava, Ariel Raskin, Elena Moreno, Sachi Johnson, Raveen Rathnasinghe, Chin I Pai, Thomas Kehrer, Elizabeth Paz Cabral, Sonia Jangra, Laura Healy, Gagandeep Singh, Prajakta Warang, Viviana Simon, Mia Emilia Sordillo, Harm van Bakel, Yonghong Liu, Weina Sun, Lisa Kerwin, Peter Palese, John Teijaro, Michael Schotsaert, Florian Krammer, Damien Bresson, Adolfo García-Sastre, Yanwen Fu, Benhur Lee, Colin Powers, Thomas Moran, Henry Ji, Domenico Tortorella, Robert Allen

## Abstract

The continual emergence of SARS-CoV-2 variants of concern, in particular the newly emerged Omicron (B.1.1.529) variant, has rendered ineffective a number of previously EUA approved SARS-CoV-2 neutralizing antibody therapies. Furthermore, even those approved antibodies with neutralizing activity against Omicron are reportedly ineffective against the subset of Omicron variants that contain a R346K substitution, demonstrating the continued need for discovery and characterization of candidate therapeutic antibodies with the breadth and potency of neutralizing activity required to treat newly diagnosed COVID-19 linked to recently emerged variants of concern. Following a campaign of antibody discovery based on the vaccination of Harbour H2L2 mice with defined SARS-CoV-2 spike domains, we have characterized the activity of a large collection of Spike-binding antibodies and identified a lead neutralizing human IgG1 LALA antibody, STI-9167. STI-9167 has potent, broad-spectrum neutralizing activity against the current SARS-COV-2 variants of concern and retained activity against the Omicron and Omicron + R346K variants in both pseudotype and live virus neutralization assays. Furthermore, STI-9167 nAb administered intranasally or intravenously provided protection against weight loss and reduced virus lung titers to levels below the limit of quantitation in Omicron-infected K18-hACE2 transgenic mice. With this established activity profile, a cGMP cell line has been developed and used to produce cGMP drug product intended for use in human clinical trials.

## INTRODUCTION

The severe acute respiratory disease syndrome coronavirus 2 (SARS-CoV-2) pandemic has continued to significantly impact the health and lives of people around the globe ^1^. To date, public health agencies have sought to combat infections leading to COVID-19 by relying on quarantine, social distancing, vaccination, and antiviral countermeasure strategies ^2, 3^. Despite these efforts, the continued spread of SARS-CoV-2 has led to the emergence of several variants of concern (VOCs) that have risen in prevalence worldwide ^2–7^.

Each VOC encodes multiple changes in the amino acid sequence of the SARS-CoV-2 spike that can impact the neutralizing properties of manufactured SARS-CoV-2 neutralizing antibodies (nAbs) as well as nAbs elicited following vaccination or during the course of natural infection. Specifically, the Omicron VOC (B.1.1.529) live virus, when profiled in vitro using Vero cells expressing human ACE2 and human TMPRSS2 for susceptibility to nAbs currently authorized or approved for clinical use (AFCU nAbs), has been shown to be resistant to the neutralizing activities of REGN10987 (imdevimab), REGN10933 (casirivimab), LY-CoV555 (bamlanivimab), LY-CoV016 (*e*tesevimab), and CT-P59 (regdanvimab), at nAb concentrations ≤ 10 μg/mL (IC_50_), and remained susceptible to nAbs COV2-2130 (cilgavimab) and COV2-2196 (tixagevimab) tested as single nAb therapies or in combination (IC_50_ of 43, 126, and 181 ng/mL, respectively) ^8–13^. In live virus neutralization assays utilizing Vero cells overexpressing human TMPRSS2, S309 (sotrovimab) registered an IC_50_ of 373 ng/mL, consistent with previously published activity in Omicron pseudovirus assays for this antibody.

A subset (approximately 23% of Omicron sequences in GISAID recorded on outbreak.info) of Omicron viruses encode an additional mutation in the SARS-CoV-2 spike at position R346K in the receptor binding domain (RBD) of the protein ^14–16^. The R346K mutation was previously identified among the defining mutations of the SARS-CoV-2 Mu VOC ^7^. Using Omicron + R346K pseudoviruses, neutralization potency was reported as substantially reduced for all tested AFCU nAbs, including COV2-2130, COV2-2196, and S309 ^5, 17–20^. Current antibodies in development, including bebtelovimab and BRII-198 (romlusevimab), maintain activity in Omicron pseudotype neutralization assays ^12, 21^. BRII-198 displays substantially reduced neutralizing activity in assays using Omicron + R346K pseudoviruses while testing of bebtelovimab against the Omicron + R346K variant has not yet been reported ^12, 21^. As such, there is a continued need for discovery and development of nAbs that can provide potent immune protection against COVID-19 caused by pandemic VOCs presently infecting the global population.

In the early COVID-19 disease setting, intravenous (IV) administration of nAbs is an effective means of lessening progression and overall severity of disease ^18, 22^. As COVID-19 is a predominantly respiratory disease, exploration of alternative modes of antibody administration including intranasal (IN) delivery may provide an expedient means of delivering antibodies and increasing the respiratory tract bioavailability of anti-COVID-19 nAbs as well as augmenting the developing host-directed immune response to prevent exacerbation of clinical symptoms and hospitalization ^23–25^.

Data presented herein demonstrate the identification, *in vitro* binding, and potent neutralizing activity of STI-9167 against live viruses and pseudotype viruses representing the current catalog of SARS-CoV-2 variants, including the Omicron and Omicron +R346K variant. Additionally, we describe the protective effects of STI-9167 administered IV or IN in the K18ACE2 transgenic mouse model of COVID-19 disease following challenge with either the WA-1 strain, Delta, or Omicron VOC.

## RESULTS

### Generation of human anti-SARS-CoV-2 spike antibodies

To generate a panel of neutralizing human monoclonal antibodies against SARS-CoV-2, Harbour *H2L2*® mice were immunized and boosted with a receptor binding domain (RBD) fusion protein based on the original spike glycoprotein sequence from the Wuhan seafood market pneumonia virus isolate (GenBank Accession# MN908947) which was fused to a mouse Fc domain (**Figure 1A**). The sera from immunized mice were assessed for binding to 293ExpiF cells transfected with SARS-CoV-2 spike cDNA (original Wuhan strain) using high-throughput flow cytometry (**Figure 1B**). We observed that the serum from Mouse 1, 3, and 4 demonstrated a concentration dependent and specific binding to 293ExpiF cells expressing SARS-CoV-2 spike. Given that Mouse 3 and Mouse 4 had the highest titer humoral response against SARS-CoV-2 spike, the spleens from these animals were used to generate hybridoma clones ^26^. The hybridoma clones (1,824 clones from Mouse 3 and 1,440 clones from Mouse 4) were screened for binding to 293ExpiF cells expressing SARS-CoV-2 spike by flow cytometry and RBD-spike (Wuhan) (**Figure 1**). A representative heat map for Mouse 4 fusion was generated to summarize the mean fluorescence intensity (MFI) for each hybridoma clone (**Figure 1C**). In parallel, hybridoma clones were subjected to an RBD ELISA validating the clones that bound to SARS-CoV-2 spike. We identified 188 clones with a >5-fold MFI over untransfected cells and classified these as candidate SARS-CoV-2 binding antibodies. The supernatants from these clones were then evaluated in a high-throughput neutralization assay using the replication competent VsV reporter virus that utilizes SARS-CoV-2 spike (VsV^CoV2-spike^) as its envelope protein and expressing GFP as readout for infection ^27^ (**Figure 1C**, Secondary Screening). Briefly, VsV^CoV2-spike^ was preincubated with hybridoma supernatant (1:20) followed by infection of HEK-293 cells expressing TMPRSS2 for 24hrs and analyzed for GFP positive cells using flow cytometry. The % infection was determined by assigning 100% infection with VsV^CoV2-spike^ pre-incubated with hybridoma media alone. Thirty-eight clones that decreased infection >50% were selected for further evaluation for neutralization by determining the IC_50_ values. For the hybridomas from the Mouse 3 fusion, 340 clones were found to bind to SARS-CoV-2 spike and 90 clones were found to have neutralization activity against VsV^CoV2-spike^. The selected neutralizing clones with IC_50_ values <125pM were then examined for IgG isotypes and expanded for further analysis. Sequencing of the heavy chain from each hybridoma clone revealed diverse CDR3 lengths ranging from 10-20 aa in length (**Figure 1D**). Clones that were identical copies of each other were consolidated to a single candidate.

**Figure 1.**
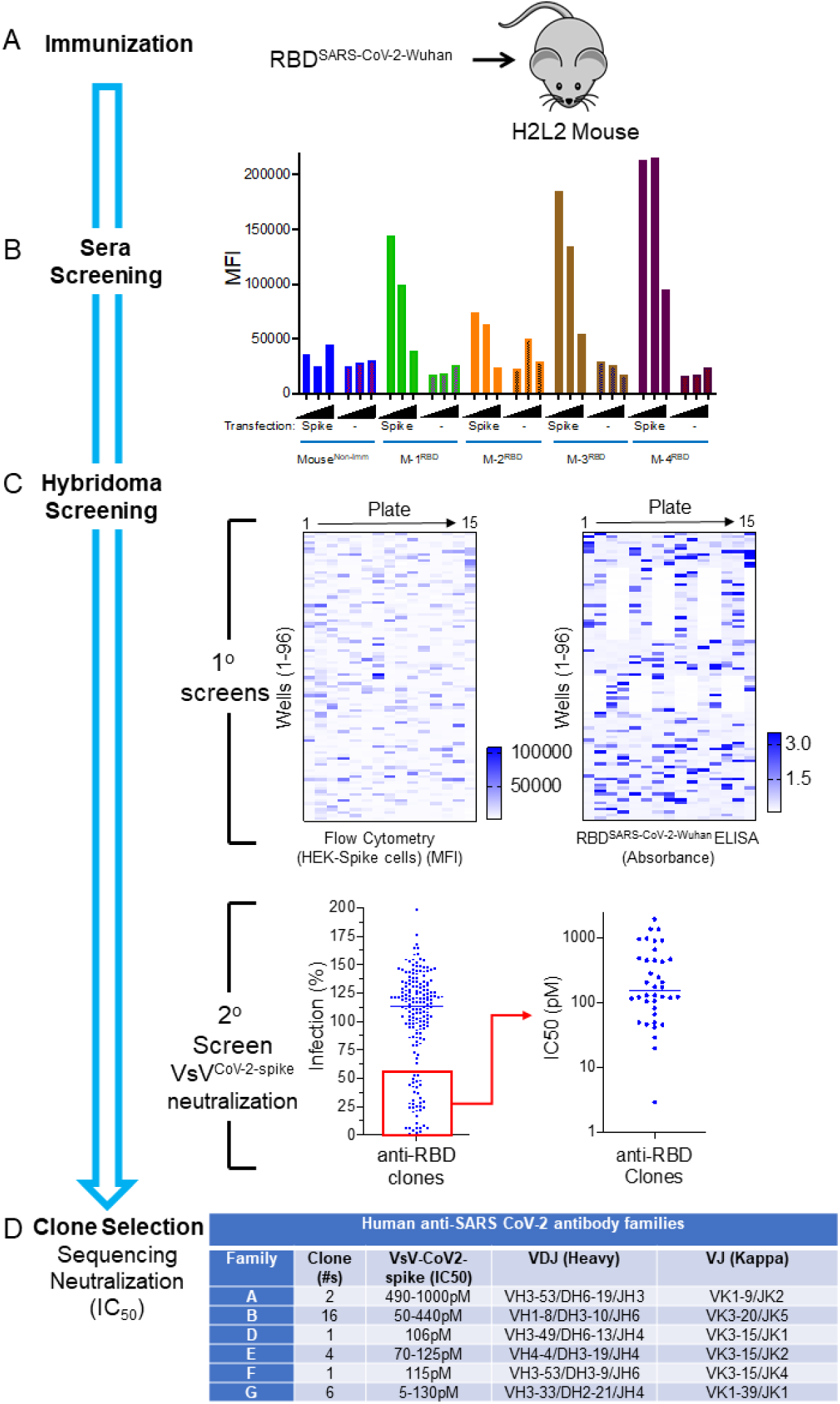
Rapid Discovery of Neutralizing Antibodies. (A) Harbour H2L2 Mice® (M-1, -2, -3, -4) were immunized and boosted 2X with SARS-CoV-2 RBD (Wuhan strain) and (B) sera (1:100, 1:500, and 1:2,500) from these mice were analyzed by flow cytometry from Expi293F untransfected or transfected with SARS-Cov-2 spike. As a control, serum from a non-immunized mouse was used. (C) A primary screen based on the anti-RBD clones from mouse 4(M-4) was performed using flow-cytometry using HEK-293 cells transfected with spike protein and RBD ELISA. Upon flow cytometry analysis, the mean fluorescence intensity (MFI) was determined for each clone. The RBD-ELISA represents binding of the clones to RBD as measured by absorbance. Both the flow cytometry and ELISA data are represented as heat maps. The secondary assay for the binding clones was a neutralization assay using VSV-spike^CoV-2^ followed by a determination of IC50 (pM) for clones with > 50% neutralization activity. (D) The clones with IC50 values <500 pM were sequenced and mAb clones were identified by specific V(D)J gene-segment combinations and junction (CDR3) characteristics, which allowed them to be grouped into different clonal families (Family “A-G”).

To identify the most effective human anti-SARS-CoV-2 spike neutralizing antibodies, we performed a VsV^CoV2-spike^ neutralization assay (**Figure 1D**). Briefly, VsV^CoV2-spike^ preincubated with increasing concentrations of antibody (0-5mg/mL) was added to TMPRSS2-expressing HEK-293 cells and analyzed for GFP positive cells using flow cytometry.

VsV^CoV2-spike^ preincubated with a control antibody was used as 100% infection. The various unique antibodies have a range of IC_50_ values from 5-1,000pM across the different antibody families. Collectively, we have identified a panel of unique human antibodies that bind to SARS-CoV-2 and effectively neutralize a reporter virus that utilizes the SARS-CoV-2 spike for entry into human cells.

Candidate nAbs sequences were formatted as full-length human IgG1 antibodies and expressed in Chinese hamster ovary (CHO) cells for further characterization in vitro. It has been shown in the context of multiple virus infections that virus-specific antibodies can lead to exacerbation of disease symptoms through a process termed antibody dependent enhancement (ADE) ^28, 29^. To reduce the risk of ADE resulting from administration of our lead candidate STI-9167, the IgG1 Fc regions were modified by introducing specific amino acid substitutions (L234A, L235A [LALA]) ^30, 31^. The LALA Fc modification reduces binding affinity to the Fcγ receptors while providing a similar blockade to interactions between SARS-COV-2 and the angiotensin-converting enzyme 2 (ACE2) receptor expressed on susceptible cells in the lung and other organs ^32–34^.

To determine the effects of variant specific spike S1 domain mutations within and outside the RBD region of S1 on antibody binding, the affinity of STI-9167 and EUA-approved SARS-CoV-2 nAbs sotrovimab, cilgavimab, and tixagevimab were determined for monomeric WA-1 spike S1 subunit binding as well as VOC-derived S1 domains using surface plasmon resonance (SPR). Of note, the k_D_ of STI-9167 was measured as 6.20 nM for the WA-1 isolate, 4.45 nM for the Delta variant, and 22.6 nM for the Omicron variant. Binding kinetics for the Omicron variant were compared between STI-9167, sotrovimab, cilgavimab, and tixagevimab. (**Figure 2A, Table 1A and Supplemental Figure 4**). STI-9167 and cilgavimab had a similar association rate and sotrovimab was approximately 5-fold slower. The dissociation rate was slowest with sotrovimab by a factor of approximately 10-fold as compared to STI-9167, and STI-9167 dissociated at an approximately 2-fold slower rate than cilgavimab. Tixagevimab binding to Omicron S1 domain monomer was insufficient to allow for quantitation.

**Table 1.** Binding and Neutralization of various Neutralizing Antibodies to SARS-CoV-2 and select VOC. (A) Omicron spike S1 binding affinity to indicated nAbs. (B) Spike protein from selected VOCs expressed on HEK 293 cells binding to presented nAbs expressed as EC50 (µg/mL). (C) Spike-pseudotyped VSV neutralization of indicated nAbs. (D) Live virus neutralization on Vero and VERO-ACE2 cells for WA-1 and Omicron virus, using indicated nAbs.

| Binding Affinity to Spike S1 Omicron |  |  |  |  |  |
| --- | --- | --- | --- | --- | --- |
| nAb | ka (1/Ms) | kd (1/s) | KD (M) | Rmax (RU) | Chi² (RU²) |
| STI-9167 | 1.69 E+05 | 4.39 E-03 | 2.60 E-08 | 231.1 | 0.51 |
| Cilgavimab | 1.02 E+05 | 1.05 E-02 | 1.03 E-07 | 42.2 | 0.58 |
| Tixagevimab | n.d. | n.d. | n.d. | n.d. | n.d. |
| Sotrovimab | 2.24 E+04 | 3.34 E-04 | 1.49 E-08 | 56.1 | 0.04 |

Abbreviation: n.d. not determined

B
| Cell Binding |  |  |  |
| --- | --- | --- | --- |
| nAb | HEK 293 cells |  |  |
|  | WA-1 | Omicron | Omicron+ R346K |
|  | EC50 (µg/mL) | EC50 (µg/mL) | EC50 (µg/mL) |
| STI-9167 | 0.014 | 0.025 | 0.025 |
| Cilgavimab | 0.084 | 0.44 | 27.2 |
| Tixagevimab | 0.0087 | 1.52 | 0.79 |
| Sotrovimab | 0.6253 | 14.5 | 2.82 |

C
| Pseudotype Neutralization |  |  |  |  |  |  |
| --- | --- | --- | --- | --- | --- | --- |
| nAb | HEK-Blue 293 hACE2-TMPRSS2 cells |  |  |  |  |  |
|  | D614G |  | Omicron |  | Omicron+R346K |  |
|  | IC50 (µg/mL) | IC80 (µg/mL) | IC50 (µg/mL) | IC80 (µg/mL) | IC50 (µg/mL) | IC80 (µg/mL) |
| STI-9167 | 0.0036 | 0.0123 | 0.0148 | 0.0774 | 0.0239 | 0.1266 |
| Cilgavimab | 0.0353 | 0.0783 | 9.105 | >10 | >10 | >10 |
| Tixagevimab | 0.0067 | 0.0184 | 0.6386 | 5.059 | 0.4696 | 3.55 |

D
| Neutralization |  |  |
| --- | --- | --- |
| nAb | Vero cells |  |
|  | SARS-CoV-2<sup>WA</sup> | SARS-CoV-2<sup>omicron</sup> |
|  | IC50 (ng/mL) | IC50 (ng/mL) |
| STI-9167 | 6.041 | 54.29 |
| Cilgavimab | 56.34 | 582.5 |
| Tixagevimab | 8.726 | 197.2 |
| Sotrovimab | 166.7 | 393 |

Abbreviation: n.t. not tested

**Figure 2.**
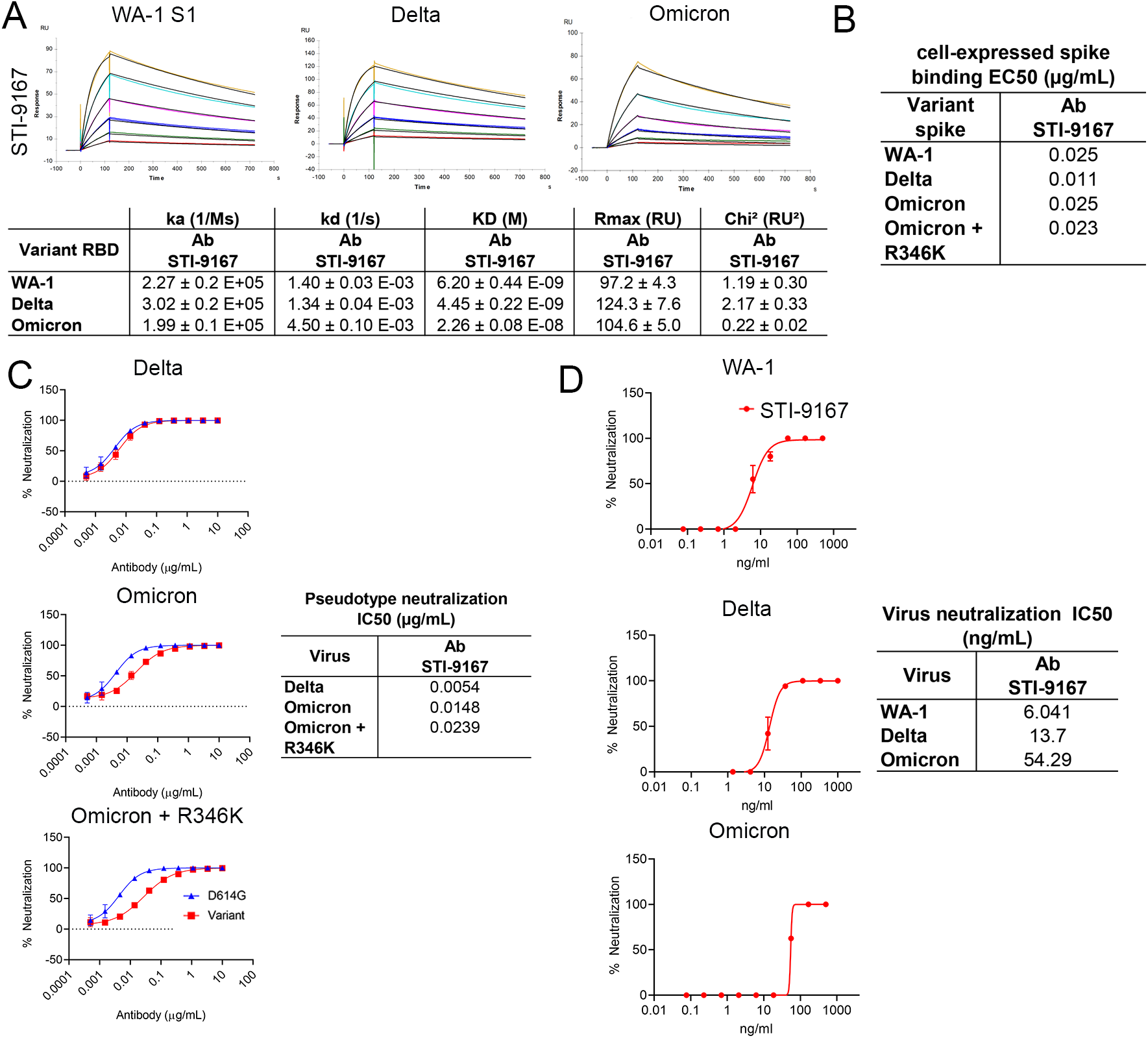
Binding and neutralization of candidate antibody. (A) Affinity measurements of STI- 9167 for Spike S1 binding domain from the following isolates and VOCs: USA/WA-1/2020(WA-1) isolate, Delta, and Omicron. The antibody affinities were measured using SPR on a BIAcore T200 instrument using a 1:1 binding model. (B) Spike protein derived from WA-1, Delta, Omicron, and Omicron + R346K SARS-CoV-2 isolates were independently expressed on the surface of HEK 293 cells. Serially-diluted STI-9167 was assayed for Spike protein binding by flow cytometry. To quantify antibody binding, mean fluorescent intensity was measured for each dilution tested and the EC_50_ value was calculated for each nAb. Representative replicate experiments are shown. (C) Spike-pseudotyped VSV neutralization. Antibody neutralization of the indicated spike variant pseudotyped VSVs was performed as described in the methods. The curves represent the average of three independent experiments, with error bars representing one standard deviation. IC50 values for each pseudotype/antibody combination are indicated on the right. (D) PRNT assay using STI-9167 with indicated SARS-COV-2 variants were performed as described in the methods, presenting percent neutralization and the calculated IC50 values indicated on the right.

In an effort to assess nAb binding to spike proteins in a native conformation, STI-9167 was tested for the binding of full-length spike protein expressed on the surface of transfected HEK293 cells. Cell-based binding studies demonstrated STI-9167 binds with similar efficiency to surface-expressed spike from the WA-1 isolate (EC_50_=0.025 µg/mL), Delta variant (EC_50_=0.011 µg/mL), and the Omicron variant (EC_50_=0.024 µg/mL), as well as the greater catalog of VOC spike protein (**Figure 2B and Supplemental Figure 1**). In general, the rank-order of binding efficiencies to surface-expressed spike for those nAbs considered in the SPR studies followed the same pattern as that determined for spike monomer binding, with the greatest concordance in binding efficiency seen between STI-9167 and cilgavimab (**Table 1B and Supplemental Figure 5**). Of note, half-maximal binding of STI-9167 to the Omicron + R346K spike (EC_50_=0.023 µg/mL) was equivalent to that measured for Omicron, suggesting that the epitope recognized by STI-9167 is preserved in the context of Omicron + R346K as compared to epitopes engaged by cilgavimab, which displayed reductions in Omicron + R346K spike binding of over 60-fold as compared to EC_50_ values in assays targeting Omicron spike. Based on the spike S1 and full-length spike protein binding data, STI-9167 was further profiled to determine the potency of virus neutralization and the breadth of neutralizing protection this antibody provided against SARS-CoV-2 variants of concern in vitro.

Virus pseudotypes were used to determine the neutralization potency (IC_50_) of STI-9167 against an index virus generated with a spike protein that carries a single D614G (VSV^D614G-spike^) mutation as compared to the WA-1 spike protein ^35^. To approximate conditions found in the setting of human SARS-CoV-2 infection, pseudovirus assays were carried out using HEK293 cells which overexpressed human ACE2 and TMPRSS2 proteins. The average IC_50_ value for STI-9167 in assays using the VSV^D614G-spike^ pseudovirus was 3.6 ng/mL (**Table 1C**). The STI-9167 neutralization potency for VSV^Delta-spike^ (IC_50_=5.4 ng/mL), VSV^Omicron-spike^ (IC_50_=14.8 ng/mL), and VSV^Omicron+R346K-spike^ (IC_50_=23.9 ng/mL) pseudotypes was maintained to within 7-fold of that measured in assays with the VSV^D614G-spike^ pseudotype (**Figure 2C**). Furthermore, STI-9167 neutralization potency was maintained to the same degree against the full catalog of VOC-based pseudovirus tested, including Alpha, Beta, Gamma, Delta Plus, Epsilon, Zeta, Iota, Kappa, Lambda, and Mu VOCs (**Supplemental Data Figure 1**). The concentration of nAbs required to achieve half-maximal and eighty-percent-maximal levels of neutralization potency for VOC pseudotypes as well as for the VSV^D614G-spike^ pseudotype are detailed in Table 1C.

The potency of STI-9167, cilgavimab, and tixagevimab was further characterized in live virus neutralization assays utilizing Vero cells. Neutralizing activity was determined following infection with WA-1 strain or Omicron variant and compared to EUA approved antibodies (**Table 1D and Supplemental Figure 3**). In keeping with the results from pseudovirus assays, we observed that STI-9167 neutralized all isolates tested including the Delta variant and Omicron variant at half-maximal concentrations within 9-fold of those measured against live WA-1 virus, with an IC_50_ of 54.29 ng/ml against live Omicron variant virus (**Figure 2D**). Neutralization potency for Omicron virus in experiments using Vero target cells was 582.5 ng/ml for cilgavimab, 197.2 ng/ml for tixagevimab, and 393 ng/ml for sotrovimab.

### Bioavailability

The biodistribution of STI-9167 was evaluated following delivery by either the intravenous or intranasal route. These studies illustrate the potential effects of delivery route on the timing of antibody exposure in the lung tissue and sera of treated mice. Following IV treatment at a dose level of 0.5 mg/kg, STI-9167 was detected in the serum, spleen, lungs, small intestine, and large intestine of most animals. Detected levels in the serum following IV dosing at the 0.5 mg/kg dose averaged 6.2 µg/mL, while STI-9167 was undetected in lung lavage material at each of the IV doses tested, (**Figure 3A**, upper left panel). Upon processing of lung tissue, antibody was detected at a mean concentration of 0.4 ng/mg of tissue in the 0.5 mg/kg IV dose group. Lower IV doses of STI-9167 did not lead to a statistically significant difference in antibody detected in lung tissue as compared to untreated animals. Antibody levels in the spleen reached an average concentration of 0.2 ng/mg of tissue within 24 hours of IV dosing at 0.5 mg/kg. Similarly, antibody was detectable in a majority of both the small and large intestines only at the highest dose level, with average concentrations of 0.14 and 0.07 ng/mg of tissue, respectively (**Figure 3A**, upper right panel).

**Figure 3.**
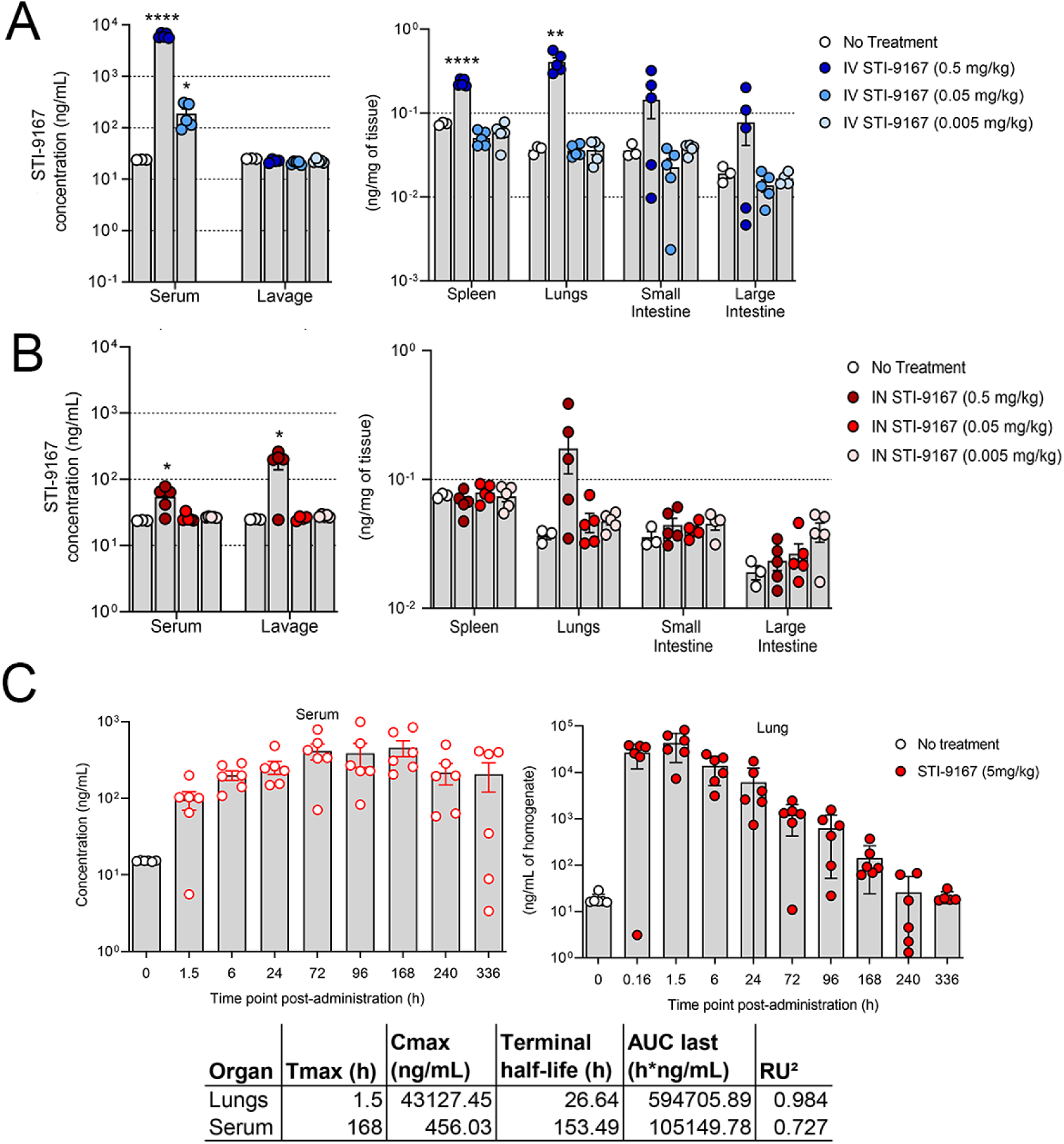
Pharmacokinetic and bioavailability of Neutralizing Antibody. *Biodistribution:* Concentration of STI-9167 in serum and lung lavage or lysates of spleens, lungs, small intestines, and large intestines collected from female CD-1 mice administered STI-9167 (A) IV at doses of 0.5 mg/kg (l), 0.05 mg/kg (l), or 0.005 mg/kg (l) or (B) IN at doses of 0.5 mg/kg (l), 0.05 mg/kg (λ), and 0.005 mg/kg (l) at 24 hours post-administration as compared to samples collected from untreated mice. Values represent mean ± SEM (n=3-4 animals no treatment group, n=5 in treatment groups). Significant differences are denoted by *, P < 0.05; **, P < 0.01; ***, P < 0.001, ****, P < 0.0001. *Pharmacokinetics:* Concentration of STI-9167 (C) in lungs and isolated serum collected from female CD-1 mice administered STI-9167 intranasally (IN) at a dose of 5 mg/kg. Samples from treated mice were collected at the indicated timepoint post-administration; antibodies concentrations were quantified by ELISA and compared to samples collected from untreated mice. Values represent mean ± SD (n=3-6 animals no treatment group, n=6 per time point in treatment groups).

Following intranasal (IN) administration of STI-9167, the concentration of antibody in the serum at 24 hours reached an average value of 0.054 µg/mL in the 0.5 mg/kg dose group. As compared to IV treated animals at the 0.5 mg/kg dose, STI-9167 administered IN resulted in a 114-fold lower concentration of antibody in serum at the 24-hour timepoint. In contrast to the observed reductions in IN serum nAb levels vs. those following IV nAb administration, STI-9167 concentrations in lung lavage samples following IN dosing reached average concentrations of 0.18 µg/mL in the 0.5 mg/kg group, a 9-fold increase over lung lavage nAb levels observed following IV delivery of the 0.5 mg/kg dose, confirming that lung lavage materials can more efficiently collect drugs delivered through the airway than those delivered IV. In lung tissue samples 24 hours following the 0.5 mg/kg IN dose, STI-9167 was detected at an average concentration of 0.173 ng/mg of tissue, similar to those levels recorded in IV-treated animals at the same dose level. STI-9167 levels in spleen, small and large intestine at all IN dose levels tested did not rise to concentrations above background (**Figure 3B**).

Overall, IN delivery of STI-9167 led to lower serum concentrations, increased lung lavage concentrations, and similar tissue concentrations in the spleen, lungs, small intestine, and large intestine when compared to IV delivery. This suggests that IN administration serves to increase the amount of antibody in the pulmonary lavage material, potentially allowing for more efficient neutralization of respiratory virus particles present in the extravascular spaces along the respiratory tract during the initial stages of infection.

### Pharmacokinetics

To characterize STI-9167 pharmacokinetic parameters following intranasal dosing, antibody levels in CD-1 mouse lung tissue lysates and serum were quantified at designated timepoints spanning a total of 336 h using a human antibody detection ELISA assay. In this assay the background concentration was on average 16.8 ng/mL based on measurements obtained using pre-dose samples. Following IN administration of STI-9167, the antibody concentration was quantifiable for most of the animals at the 336 h timepoint in both the lungs and serum (**Figure 3C**). The C_max_ value of STI-9167 in the lungs was measured at 1.5 hours (T_max_) post-administration at a value of 43 μg/mL. In the lungs following IN administration, STI-9167 exhibited an apparent terminal half-life (T_1/2_) of 26.6 h. Kinetics of STI-9167 exposure in the lungs following IN administration contrasted with the slower rate of antibody accumulation in the serum of treated mice (**Figure 3C**). Antibody was first detected in the serum at 1.5 hours post-administration and the C_max_ of 456 ng/mL was reached at the 168 h timepoint (T_max_), although consideration of the standard deviations in values measured among animals on or between the 72h and 168h timepoints suggests that the T_max_ may have occurred as early as 72 hours post-administration. Antibody levels remained relatively constant in serum over the period spanning 24 - 336 h, which is in keeping with the calculated STI-9167 serum half-life observed following IV STI-9167 administration in CD-1 mice (data not shown). The total systemic antibody exposure (AUC_last_) was greater than 5-fold higher in the lungs than in the serum of IN-treated mice (AUC_last_ were 594,705 and 105,149 h*ng/mL respectively).

### Treatment using IV or IN administered STI-9167 in the K18-hACE2 transgenic mouse model of COVID-19

SARS-CoV-2 pathogenesis in the K18-ACE2 transgenic model of COVID-19 respiratory disease provides a tractable means of assessing nAb activity in a preclinical model of respiratory disease ^36–38^. The clinical signs and histological markers of pathogenesis in this model include weight loss over the first four to five days post-infection and the presence of microscopic lesions in the infected lungs ^37–40^. Peak virus lung titers are typically detected by day 5 post-infection, but the timing and peak amplitude of replication in the lungs can vary depending on the specific VOC used to challenge the mice ^39^. The breadth of protection provided by STI-9167 was established by treating mice following virus challenge with 1 x 10^5^ half-maximal tissue culture infectious dose (TCID_50_) of the WA-1 strain, the Delta variant, or the Omicron variant. Animals treated with isotype control antibody lost weight in each experiment, with the average percentage weight reduced to 91.3 % with WA-1 strain, 89 % with Delta variant, and 94.7 % with Omicron variant (**Figure 4B, 4E**) as compared to average Day 0 weights in each group. To determine the effects of the route of administration on the degree of protection conferred by treatment with STI-9167 at doses ranging from 5 to 20 mg/kg, antibody was administered by either intravenous injection (**Figure 4A**) or intranasal instillation (**Figure 4D**). At a dose level of 5 mg/kg, administration of STI-9167 to K18-hACE2 mice by either the IV or IN route provided protection against weight loss caused by WA-1 strain, Delta variant, and Omicron variant (**Figure 4B, 4E**). Virus replication in the lungs, quantified on day 4 post-infection, was approximately 2.5 x 10^6^, 2.1 x10^4^, and 4.5 x 10^2^ TCID_50_/g on average in isotype control-treated mice infected with WA-1, Delta variant, or Omicron variant, respectively. Following infection by each of the SARS-CoV-2 challenge viruses, lung virus titers in mice treated with STI-9167 were reduced to levels below the limit of quantification, independent of the nAb dosing route or the nAb dose level (**Figure 4C, 4F**).

**Figure 4.**
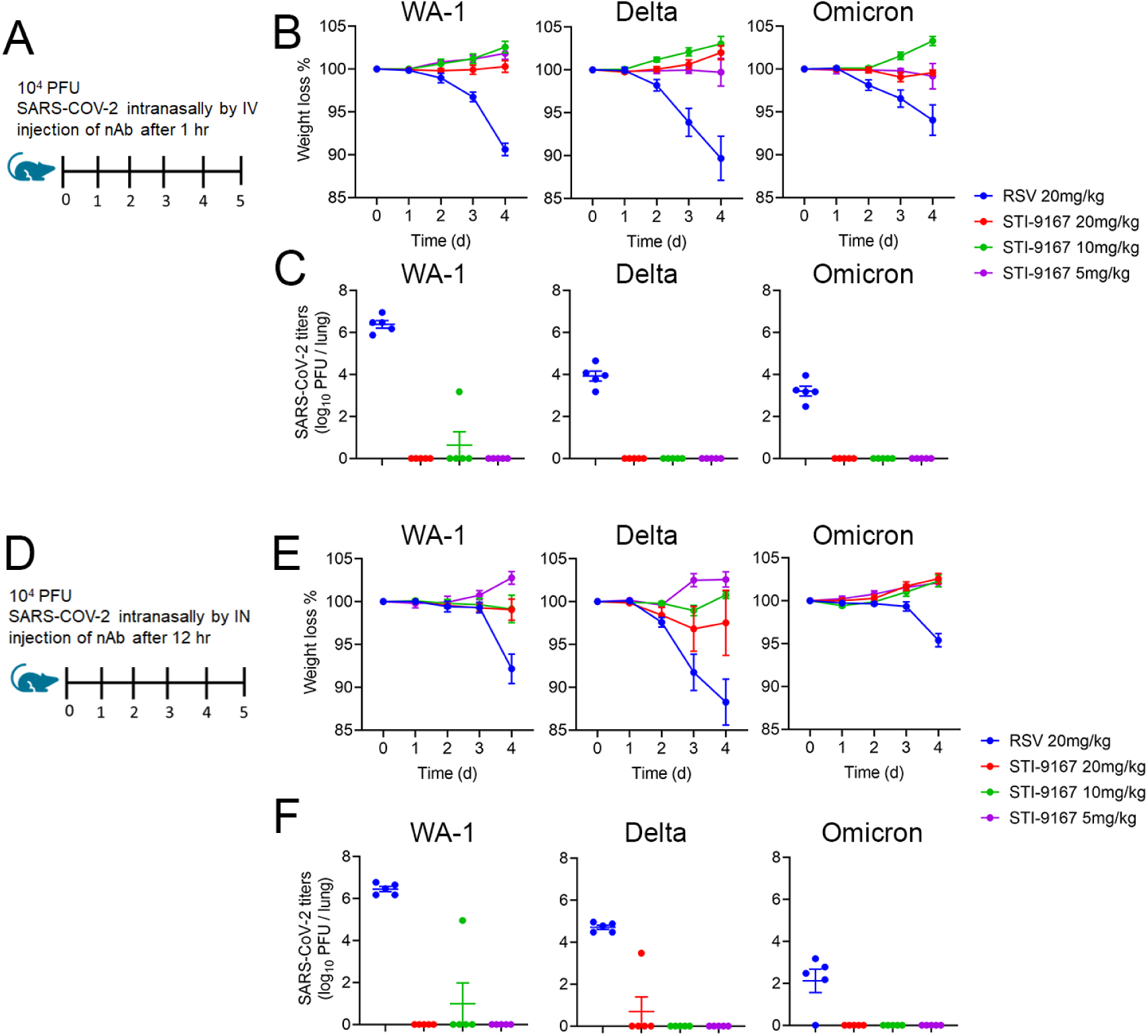
Efficacy of Intranasal (IN) delivery of STI-9167 Neutralizing Antibody in the K18-hACE2 murine model of COVID-19. (A) K18-hACE2 transgenic mice were infected with 10,000 PFU of WA-1, Delta or Omicron SARS-CoV-2 treated with indicated concentration of isotype control antibody (Isotype) or STI-9167 intravenously 1 h post infection. (B) Body weight change of mice was measured daily (n = 5). (C) SARS-CoV-2 viral titers were measured in lung day 4 post infection (n = 5). (D) K18-hACE2 transgenic mice were infected with 10,000 PFU SARS-CoV-2 WA1, Delta, or Omicron strains and treated with indicated concentration of Isotype or STI-9167 intranasally 12 h post infection.(E) Body weight change of mice was measured daily (n = 5). (F) SARS-CoV-2 viral titers were measured in lung day 4 post infection (n = 5) n.s. not significant, P* < 0.05, P** < 0.01, P*** < 0.001 or P****<0.0001. Unpaired t-test (C and F). Two way ANOVA (B and E)

## DISCUSSION

Use of antibody discovery platforms that do not require material derived from infected individuals, such as the vaccination strategy employed here or the screening of established antibody libraries, can provide a preemptive means of addressing the challenges presented by pandemic threat pathogens ^14, 41^. The production of human antibodies in transgenic animals has several advantages including *in vivo* affinity maturation, increased diversity, and clonal selection for antibody optimization ^42^. Thus, the generation of antibodies to specific protein domains allows for the development of highly reactive and effective antibody therapeutics.

The pool of antibodies we identified following vaccination of mice with an SARS-CoV-2 RBD protein based on the Wuhan spike protein sequence includes a candidate with potent neutralizing activity against many SARS-CoV-2 variants of concern that have emerged in the past two years of the pandemic. Our antibody binding studies and virus neutralization assays have provided clear evidence of the broad and potent neutralizing activity of STI-9167 toward those VOCs identified in the early period of the pandemic as well as those VOCs currently impacting public health, including Delta and Omicron. Following demonstration of neutralizing activity against the parental Omicron variant, we have extended our activity profiling studies to include assessment of the Omicron + R346K subvariant, a virus that is currently represented in a reported 23% of sequences submitted to GISAID ^14–16^. The frequency of Omicron virus sequences containing the R346K substitution has risen steadily since the first reports in November of 2021 describing detection of the Omicron variant ^43–45^. Using virus pseudotypes, we have demonstrated durable STI-9167 activity against the Omicron +R346K subvariant. In addition, we described neutralizing activity against the Mu variant, which also encodes the R346K spike substitution ^7^. The Omicron and Mu variants constitute divergent variants of SARS-CoV-2, and it appears that the R346K substitution is not sufficient in either of these contexts to provide a means of resistance to the neutralizing effects of STI-9167.

Our studies of nAb STI-2020 previously demonstrated the protective efficacy of IN-administered antibodies in the context of SARS-CoV-2 preclinical models of pathogenesis ^46^. Previous work in preclinical models of respiratory virus pathogenesis support the use of IN-administered IgG and IgA mAbs in prophylactic and therapeutic dosing regimens ^24, 47–50^. In the current report, we described the protective effects of STI-9167 delivered by either the IN or the IV route to animals infected with WA-1 strain, Delta, or Omicron variants. As evidenced in recently reported preclinical studies of Omicron pathogenesis as well as our experiments, the severity of clinical signs and the amount of virus replication in the lungs following Omicron infection was reduced in comparison to that following infection with the WA-1 strain or the Delta variant ^51–53^. Independent of the challenge virus used, at a dose level of 5 mg/kg, IN treatment with STI-9167 in K18 ACE2 transgenic mice 12 hours following infection provided protection against the weight loss observed in control animals and also reduced virus lung titers to below the level of quantitation. Phase 1 clinical studies with STI-2099 (plutavimab) have demonstrated the safety of nAb delivered as formulated liquid drops to the upper airways. A Phase 2 study has completed enrollment in the US, and additional Phase 2 studies are ongoing in Mexico and the United Kingdom. Based on the favorable in vivo potency and physicochemical profile of STI-9167, cGMP drug product has been prepared in preparation for similar anticipated clinical studies of STI-9167 administered IV or as intranasal drops (STI-9199).

## MATERIALS AND METHODS

### Immunizations and Hybridoma Generation

To generate human antibodies, Harbour *H2L2*® human antibody transgenic mice (Harbour BioMed, Cambridge, MA) were utilized under a collaboration between the Icahn School of Medicine at Mount Sinai and Harbour BioMed. The H2L2 transgenic mouse is a chimeric transgenic mouse containing the human variable gene segment loci of the heavy and kappa antibody chains along with the rat heavy and kappa constant gene segment loci, producing a mouse with normal B cell homeostasis and effector functions, while also producing antibodies that represent the typical diversity observed in human antibody immunity ^54^. Immunizations were done on eight-to twelve-week-old H2L2 mice interperitoneally with 50-100 μg of a recombinant SARS-CoV2 Spike RBD_319-591_-Fc fusion protein generated from sequence from the original Wuhan seafood market pneumonia virus isolate (GenBank Accession# MN908947) and cloned in-frame into pcDNA vectors containing human IgG1 and mouse IgG2a Fc tags (GenScript USA Inc., Piscataway, NJ). Each mouse received a prime followed by 2 boosts, and blood was collected from the submandibular vein two weeks after each boost to monitor titer of sera antibodies. Following sera binding and neutralization analysis, two mice were selected for hybridoma fusion and received two final boosts consisting of 50-100 μg of the RBD protein at - 5 and -2 days before being euthanized by IACUC approved methods with spleens harvested and a final bleeding collected for sera analysis (“fusion sera”). The spleens were processed to single cell suspension and hybridomas were generated using the standard protocol. Briefly, individual B cell clones were grown on soft agar and selected for screening using a robotic ClonaCell Easy Pick instrument (Hamilton/Stem Cell Technology). Individual clones were expanded, and the supernatants were used to screen for binding, neutralization and ACE2 competition assays. All animal studies were approved by the Icahn School of Medicine Institutional Animal Care and Use Committee (IACUC).

Hybridoma Screening: Expi293F cells were transiently transfected to express SARS-CoV-2 spike (Wuhan) using Lipofectamine 3000 (L3000001, Thermo Fisher) and then incubated with supernatant from the hybridoma cell lines from each fusion. Binding was detected using an anti-rat IgG-APC detection antibody and samples were run on a high-throughput flow cytometer (Intellicyte High Throughput Flow Cytometer [Intellicyte Corp., Albuquerque, NM]. Samples were compared to controls of fusion sera, unimmunized “normal” mouse sera, and an in-house generated anti-SARS1&2 Spike mouse monoclonal 2B3E5 at 1 μg/mL. Cells with a high mean fluorescence intensity (MFI) were identified using FlowJo software (Tree Star, Inc.) and graphed using GraphPad Prism to create a heat map based on mean fluorescence intensity.

ELISA: Immulon 4 HBX high binding clear flat bottom 96 well plates (ThermoFisher) were coated with SARS-CoV2 Spike RBD_319-591_-Fc fusion at 5 μg/ml in 1xPBS overnight at 4 °C followed by washing. Washing with 1xPBS was done between each step in triplicate using a Biotek ELX405 MultiPlate Washer (Biotek, Winooski, VT). Plates were blocked for two hours in blocking solution (1xPBS, 0.5% BSA). Supernatants from the hybridomas were then added and allowed to incubate for one hour at room temperature followed by the addition of goat anti-rat IgG (heavy chain specific)-HRP (Jackson ImmunoResearch) at a 1:5,000 dilution in blocking solution for one hour. ABTS substrate solution (ThermoFisher) was added and allowed to incubate for 5-10 minutes at room temperature, protected from light. Absorbance at 450nm was measured using a Biotek Synergy HT Microplate Reader. Fusion sera, normal mouse sera, and 2B3E5 mAb (0.5-1 μg/ml) were used as controls.

Neutralization: Prior to neutralization, hybridoma supernatants grown in SFM (sera free hybridoma media) (Invitrogen) were quantitated using an Octet Red96 by diluting supernatants 1:5 and 1:10 in sera free media and measured for binding against the Anti-Murine IgG Quantitation (AMQ) Biosensors (with cross reactivity to rat IgG Fc) on an Octet Red 96 BLI Instrument (SartoriusAG, Goettingen, Germany). Results were compared to in-lab derived purified rat IgG standards diluted in SFM in the range of 0.5-50 μg/ml. For neutralization, VsV-SARS-spike GFP-expressing reporter virus was pre-incubated with mouse sera (1:100-1,200), hybridoma supernatants (1:10-1:10,000), or purified human monoclonal antibodies (0.1 ng/ml-1 μg/ml) and incubated at 4 °C for 1 hr before the inoculum was added to HEK-293 cells expressing human ACE2 and Transmembrane Serine Protease-2 overnight at 37 °C, 5% CO_2_ ^27^. The cells were resuspended in cold FACS buffer and analyzed by flow cytometry (Intellicyte Corp., Albuquerque, NM) for GFP fluorescence intensity. Cells with a high MFI were identified using FlowJo software (Tree Star, Inc.) and graphed using GraphPad Prism to create a heat map based on MFI.

### Sequencing and Humanizing of Antibodies

Sequence of the human variable heavy and kappa chains were obtained by using SMARTer 5’ RACE technology (Takara Bio USA) adapted for antibodies to amplify the variable genes from heavy and kappa chains for each hybridoma. Briefly, RNA was extracted from each hybridoma using Qiagen RNeasy Mini Kit (Qiagen, Valencia, CA), followed by first stand cDNA synthesis using constant gene specific 3’ primers based on the specific isotype of the hybridoma and incubation with the SMARTer II A Oligonucleotide and SMARTscribe reverse transcriptase. Amplifying PCR of the first stand cDNA product was then performed using SeqAmp DNA Polymerase (Takara) with a nested 3’ primer to the constant genes and a 5’ universal primer based on universal primer sites added to the 5’ end during cDNA generation. Purified PCR product was then submitted for Sanger sequencing using 3’ constant gene primers (GeneWiz, South Plainfield, NJ). Sequence results were blasted against the IMGT human databank of germline genes using V-Quest (http://imgt.org) and analyzed for clonality based on CDR3/junction identity and V(D)J usage. Unique clones were chosen from each clonal family, and DNA was synthesized and cloned in-framed into pcDNA-based vectors containing a human IgG1 constant region and a human kappa light chain constant region (GenScript USA Inc., Piscataway, NJ).

### Synthesis of Comparison Antibodies

Antibody expression vector construction and antibody transient expression and purification were done following standard protocols. Briefly, heavy chain and light chain variable domain genes were designed by coding the amino acid sequences of an antibody using codon table of *Cricetulus griseus* for CHO (Chinese hamster ovarian) cells as expression host. The heavy chain and light chain variable domain gene fragments with flanking sequences for infusion cloning were synthesized by IDT (Integrated DNA technologies, San Diego), and cloned into a mammalian expression vector with built-in IgG1 constant domain and/or light chain constant domain sequences. The expression vectors were confirmed by DNA sequencing. CHO-S cells in exponential phase at a cell density of 10^6^ cells/ml with viability of ≥ 93%, were co-transfected with both heavy and light chain expression plasmids of the target antibody. The transfection complex was formed between DNA and PEI (polyethylenimine). Each antibody was harvested by centrifuging the culture to pellet and remove the cells 10-14 days after the transfection. The supernatant was processed with a protein A column, and the Protein A bound antibody was eluded with low pH glycine buffer. Purity of the antibodies were annualized by SDSPAGE to be more than 95%.

Reference for antibody sequences is as follows; Sotrovimab (https://www.kegg.jp/entry/D12014) Cilgavimab and Tixagevimab (https://www.genome.jp/entry/D11993) (https://www.genome.jp/entry/D11994)Antibody characterization

Kinetic interactions between the antibodies and his-tagged antigen proteins were measured at room temperature using Biacore T200 surface plasmon resonance (GE Healthcare). Anti-human fragment crystallizable region (Fc region) antibody was immobilized on a CM5 sensor chip to approximately 8,000 resonance units (RU) using standard N-hydroxysuccinimide/N-Ethyl-N′-(3- dimethylaminopropyl) carbodiimide hydrochloride (NHS/EDC) coupling methodology. The antibody (1.5 μg/mL) was captured for 60 seconds at a flow rate of 10 μL/minute. The SARS-CoV-2 Spike S1, SARS-CoV-2 (2019-nCoV) Spike S1-B.1.1.7 lineage mut (HV69-70 deletion, Y144 deletion, N501Y, A570D, D614G, P681H)-His and SARS-CoV-2 (2019-nCoV) Spike S1-B.1.351 lineage mut (K417N, E484K, N501Y, D614G)-His proteins were run at six different dilutions in a running buffer of 0.01 M HEPES pH 7.4, 0.15 M NaCl, 3 mM EDTA, 0.05% v/v Surfactant P20 (HBS-EP+). All measurements were conducted in HBS-EP+ buffer with a flow rate of 30 μL/minute. The affinity of antibody was analyzed with BIAcore T200 Evaluation software 3.1. A 1:1 (Langmuir) binding model is used to fit the data.

### Cell based Spike binding assay

Mammalian expression vectors were constructed by cloning of the synthesized gene fragments encoding SARS-CoV-2 Spike variant proteins (see attached table indicating mutations introduced into wild type [WA-1 strain] spike protein sequence). HEK293 cells were transfected using FuGeneHD transfection reagent according to manufacturer’s protocol (Promega, Cat # E2311). 48 hours post-transfection, cells were harvested using enzyme free cell dissociation buffer (ThermoFisher, Cat #13151014.), washed once and resuspended in FACS buffer (DPBS + 2% FBS) at 2×10^6^ cells/mL. For antibody binding to the cells expressing the Spike proteins, the cells were dispensed into wells of a 96-well V bottom plate (40 µL per well), and an equal volume of 2x final concentration of serially-diluted anti-S1 antibody solution was added. After incubation on ice for 45 minutes, the cells were washed with 2 times of 150 µL FACS buffer. Detection of bound antibody was carried out by staining the cells with 50 µL of 1:500 diluted APC AffiniPure F(ab’)₂ Fragment (Goat Anti-Human IgG (H+L). Jackson ImmunoResearch, Cat# 109-136-4098) for 20 minutes on ice. The cells were washed once with 150 µL FACS buffer and analyzed on IntelliCyt iQue® Screener (Sartorius) flow cytometry. Mean fluorescent intensity values were obtained from the histograms. A sigmoidal four-parameter logistic equation was used for fitting the MFI vs. mAb concentration data set to extract EC50 values (GraphPad Prism 8.3.0 software).

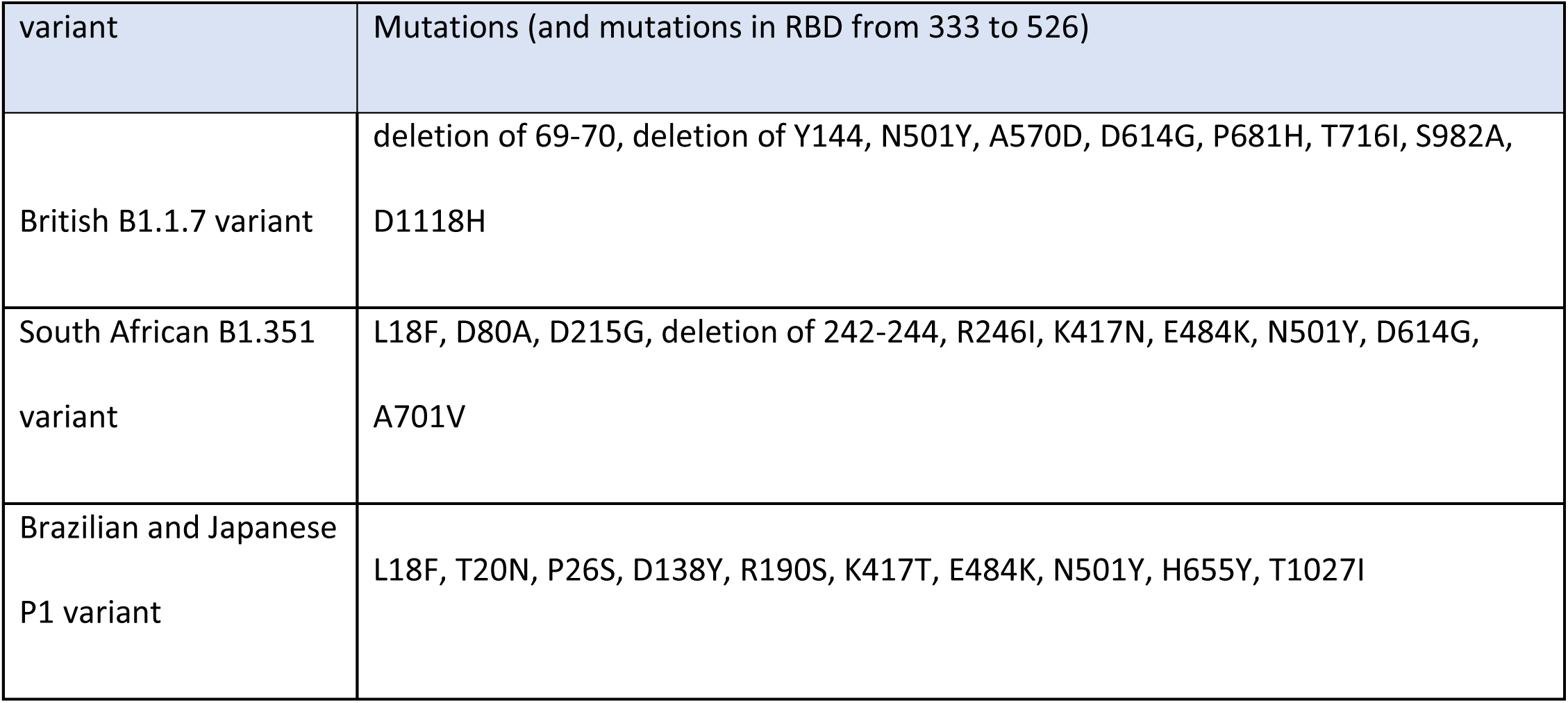
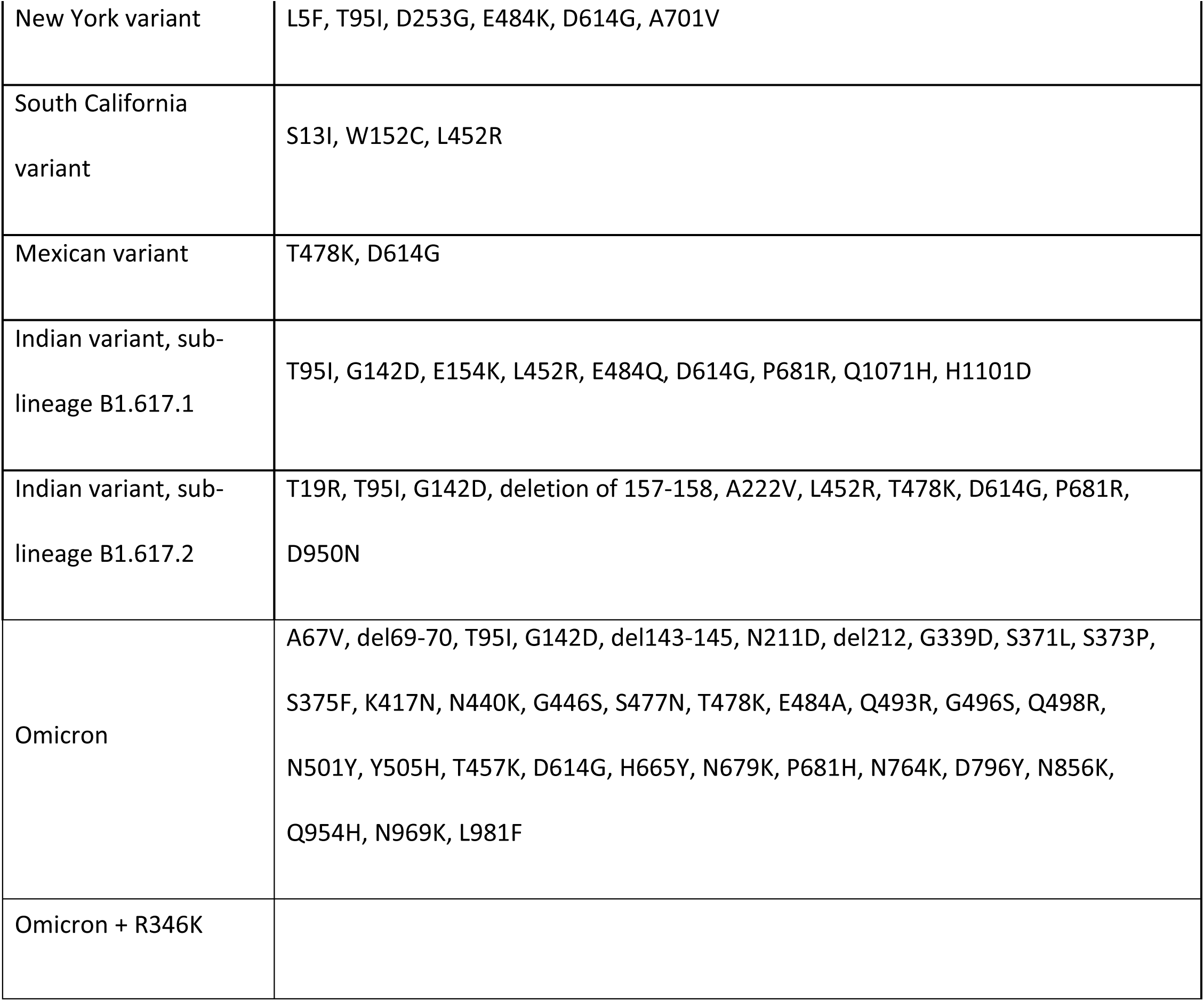

| variant | Mutations (and mutations in RBD from 333 to 526) |
| --- | --- |
| British B1.1.7 variant | deletion of 69-70, deletion of Y144, N501Y, A570D, D614G, P681H, T716I, S982A, D1118H |
| South African B1.351 variant | L18F, D80A, D215G, deletion of 242-244, R246I, K417N, E484K, N501Y, D614G, A701V |
| Brazilian and Japanese P1 variant | L18F, T20N, P26S, D138Y, R190S, K417T, E484K, N501Y, H655Y, T1027I |
| New York variant | L5F, T95I, D253G, E484K, D614G, A701V |
| South California variant | S13I, W152C, L452R |
| Mexican variant | T478K, D614G |
| Indian variant, sub-lineage B1.617.1 | T95I, G142D, E154K, L452R, E484Q, D614G, P681R, Q1071H, H1101D |
| Indian variant, sub-lineage B1.617.2 | T19R, T95I, G142D, deletion of 157-158, A222V, L452R, T478K, D614G, P681R, D950N |
| Omicron | A67V, del69-70, T95I, G142D, del143-145, N211D, del212, G339D, S371L, S373P, S375F, K417N, N440K, G446S, S477N, T478K, E484A, Q493R, G496S, Q498R, N501Y, Y505H, T457K, D614G, H665Y, N679K, P681H, N764K, D796Y, N856K, Q954H, N969K, L981F |
| Omicron + R346K |  |

### Cells and Viruses

Vero E6 cells were maintained in Dulbecco’s modified Eagle’s medium (DMEM, Corning, NY) supplemented with 10% fetal bovine serum (FBS, Thermo Fisher Scientific, MA), 1% penicillin– streptomycin, and L-glutamine. The P3 stock of the SARS-CoV-2 USA/WA-1/2020, 202001, USA/CA-CDC5574/2020 and, MD-HP01542/2021 isolates were obtained from The World Reference Center for Emerging Viruses and Arboviruses (WRCEVA) at the University of Texas Medical Branch. The viruses were propagated in Vero E6 cells and cell culture supernatant of P4 stocks were stored at -80 °C under BSL3 conditions.

BHK21 cells (ATCC #CCL-10) were maintained in DMEM/F12 media (Thermo Fisher #21041025) supplemented with 10% fetal bovine serum (Omega Scientific #FB-02) and 5% tryptose phosphate broth (Thermo Fisher #18050039). BHK21/WI-2 cells (Kerafast #EH1011) were maintained in DMEM (Thermo Fisher #11965092) supplemented with 5% fetal bovine serum. 293-ACE2 cells were maintained in DMEM supplemented with 10% fetal bovine serum and 200 µg/mL G418 (Invivogen #ant-gn-2). HEK-Blue 293 hACE2-TMPRSS2 cells (Invivogen #hkb-hace2tpsa) were maintained in DMEM supplemented with 10% fetal bovine serum, 0.5 µg/mL Puromycin (Invivogen #ant-pr-1), 200 μg/mL Hygromycin-B (Invivogen #ant-hg-1), and 100 μg/mL Zeocin (Invivogen #ant-zn-1).

SARS-COV-2 viruses were obtained from BEI resources (Washington strain NR-52281; Alpha variant NR-54000; Beta Variant NR-54009; Gamma variant NR-54982; Delta variant NR-55611 or NR-55672; Lambda variant NR-55654: Omicron Variant NR-65461.) VeroE6 monolayers were infected at an MOI of 0.01 in 5 mL virus infection media (DMEM + 2% FCS +1X Pen/Strep). Tissue culture flasks were incubated at 36 °C and slowly shaken every 15 minutes for a 90- minute period. Cell growth media (35 mL) was added to each flask and infected cultures were incubated at 36 °C/5% CO2 for 48 hours. Media was then harvested and clarified to remove large cellular debris by room temperature centrifugation at 3,000 rpm.

### SARS-CoV-2 neutralization assay

The day before infection, 2×10^4^ Vero E6 cells were plated to 96-well plates and incubated at 37 °C, 5% CO_2_. Monoclonal antibodies were 2-fold serially diluted in infection media (DMEM+2% FBS). Sixty microliters of diluted samples were incubated with 200 µL of 50% tissue culture infective doses (TCID_50_) of SARS-CoV-2 in 60 µL for 1 h at 37 °C. One-hundred microliters of the antibody/virus mixture were subsequently used to infect monolayers of Vero E6 cells grown on 96-well plates. Cells were fixed with 10% formalin and stained with 0.25% crystal violet to visualize cytopathic effect (CPE). The neutralizing concentrations of monoclonal antibodies were determined by complete prevention of CPE.

### Plasmids

All SARS-CoV-2 Spike constructs for pseudotype generation were expressed from plasmid pCDNA3.1 (ThermoFisher #V79020). Codon optimized SARS-CoV-2 Wuhan Spike carrying the D614G amino acid change (Sino Biological #VG40589-UT(D614G)) was modified to remove the last 21 amino acids at the C-terminus (SpikeΔ21) and was used as the parental clone. Amino acid changes for each variant are as follows. Alpha: Δ69-70, Δ144, N501Y, A570D, D614G, P681H, T716I, S982A, and D1118H. Beta: D80A, D215G, Δ242-244, K417N, E484K, N501Y, D614G, and A701V. Epsilon: S13I, W152C, L452R, and D614G. Kappa: G142D, E154K, L452R, E484Q, D614G, P681R, Q1071H, and H1101D. Delta: T19R, G142D, Δ156-157, R158G, L452R, T478K, D614G, P681R, and D950N. Delta Plus: T19R, G142D, Δ156-157, R158G, K417N, L452R, T478K, D614G, P681R, and D950N. Gamma: L18F, T20N, P26S, D138Y, R190S, K417T, E484K, N501Y, D614G, H655Y, and T1027I. Zeta: E484Q, F565L, D614G, and V1176F. Lambda: G75V, T76I, R246N, Δ247-253, L452Q, F490S, D614G, and T859N. B.1.1.318: T95I, ΔY144, E484K, D614G, P681H, and D796H. Mu: T95I, Y144T, Y145S, ins146N, R346K, E484K, N501Y, D614G, P681H, and D950N. Omicron: A67V, del69-70, T95I, G142D, del143-145, N211D, del212, G339D, S371L, S373P, S375F, K417N, N440K, G446S, S477N, T478K, E484A, Q493R, G496S, Q498R, N501Y, Y505H, T457K, D614G, H665Y, N679K, P681H, N764K, D796Y, N856K, Q954H, N969K, L981F

### VSV-Spike pseudotype generation

To generate each Spike pseudotyped VSV, 1.2E6 BHK21 cells were nucleofected with 2 µg of Spike plasmid using an Amaxa Nucleofector II with cell line kit L (Lonza #VCA-1005) and program A-031. Cells were plated to one well of a 6-well dish and incubated overnight at 37 °C/5%CO_2_. The next day, cells were transduced with G-Pseudotyped ΔG-luciferase (G*ΔG- luciferase) rVSV (Kerafast #EH1025-PM) at MOI∼4 for 1 hour at 37 °C/5%CO_2_. Cells were rinsed twice with DPBS (Corning #21-031-CM), 2 mL of fresh media added, and incubated for 24-44 hours at 37 °C/5%CO_2_. Supernatants were collected, spun at 300*g* for 5 minutes at room temperature, aliquoted and stored at –80 °C. Pseudotypes were normalized for luciferase expression by incubating with 1 µg/mL anti-VSV-G clone 8G5F11 (Millipore #MABF2337) for 30 minutes at room temperature followed by transduction of 293-ACE2 cells. G*ΔG-luciferase VSV of known titer was used as the standard. Transduced cells were incubated for 24 hours, 40 µL of ONE-Glo reagent (Promega #E6110) added and luminescence measured using a Tecan Spark plate reader.

### Pseudotype virus neutralization assays

HEK-Blue 293 hACE2-TMPRSS2 cells were plated to white-walled 96-well plates at 40K cells/well and incubated at 37 °C/5% CO_2_. The next day, pseudotyped VSV was incubated with anti-spike (concentration as indicated) and anti-VSV-G (1 µg/mL) antibodies for 30 minutes at room temperature and added to the HEK-Blue 293 hACE2-TMPRSS2 cells in triplicate. Transduced cells were incubated for 24 hours, 40 µL of ONE-Glo reagent (Promega #E6110) added and luminescence measured using a Tecan Spark plate reader. The percent inhibition was calculated using 1-([luminescence of antibody treated sample]/[average luminescence of untreated samples]) x 100. Absolute IC50 was calculated using non-linear regression with constraints of 100 (top) and 0 (baseline) using GraphPad Prism software. The average of triplicate samples in each of at least 3 independent experiments were included in the analyses. Negative value slopes were assigned IC50 of >10 µg/mL. IC80 values were calculated using non-linear regression with F=80 and constraints of 100 (top) and 0 (bottom). For antibody comparison experiments, data for Omicron and Omicron+R346K is an average of 2 independent experiments.

### Affinity measurements

Kinetic interactions between the antibodies and his-tagged antigen proteins were measured at room temperature using Biacore T200 surface plasmon resonance (GE Healthcare). Anti-human fragment crystallizable region (Fc region) antibody was immobilized on a CM5 sensor chip to approximately 8,000 resonance units (RU) using standard N-hydroxysuccinimide/N-Ethyl-N′-(3- dimethylaminopropyl) carbodiimide hydrochloride (NHS/EDC) coupling methodology. The antibody (0.5-1 μg/mL) was captured for 60 seconds at a flow rate of 10 μL/minute. Each of the following five variants of SARS-CoV-2 Spike S1 proteins were run at six different dilutions in a running buffer of 0.01 M HEPES pH 7.4, 0.15 M NaCl, 3 mM EDTA, 0.05% v/v Surfactant P20 (HBS-EP+). All measurements were conducted in HBS-EP+ buffer with a flow rate of 30 μL/minute. The affinity of antibody was analyzed with BIAcore T200 Evaluation software 3.1. A 1:1 (Langmuir) binding model is used to fit the data.

1. Spike S1 (wt)
2. Spike S1 (UK): HV69-70 deletion, Y144 deletion, N501Y, A570D, D614G, P681H)
3. Spike S1 (SA): K417N, E484K, N501Y, D614G
4. Spike S1 (BZ): L18F, T20N, P26S, D138Y, R190S, K417T, E484K, N501Y, D614G, H655Y
5. Spike S1 (DT): T19R, G142D, E156G, 157-158 deletion, L452R, T478K, D614G, P681R
6. Spike S1 (Omicron): A67V, del69-70, T95I, G142D, del143-145, N211D, del212, G339D, S371L, S373P, S375F, K417N, N440K, G446S, S477N, T478K, E484A, Q493R, G496S, Q498R, N501Y, Y505H, T457K, D614G, H665Y, N679K, P681H, N764K, D796Y, N856K, Q954H, N969K, L981F

### Biodistribution Study

Female CD-1-IGS (strain code #022) were obtained from Charles River at 6-8 weeks of age. For intravenous injection of 10A3YQYK, 100 µL of antibody diluted in 1X formulation buffer C was administered retro-orbitally to anesthetized animals. For intranasal injections, antibody was diluted in 1X formulation buffer C and administered by inhalation into the nose of anesthetized animals in a total volume of 20-25 µL using a pipette tip. Organs, blood, and lung lavage samples were collected 24 hours post-antibody administration. Blood was collected by retro-orbital bleeding and then transferred to Microvette 200 Z-Gel tubes (Cat no# 20.1291, lot# 8071211, SARSTEDT). Tubes were then centrifuged at 10,000*g* for 5 minutes at room temperature. Serum was transferred into 1.5 mL tubes and stored at −80 °C. Lung lavage samples were collected following insertion of a 20G x 1-inch catheter (Angiocath Autoguard, Ref# 381702, lot# 6063946, Becton Dickinson) into the trachea. A volume of 0.8 mL of PBS was drawn into a syringe, placed into the open end of the catheter, and slowly injected and aspirated 4 times. The syringe was removed from the catheter, and the recovered lavage fluid was transferred into 1.5 mL tubes and kept on ice. Lavage samples were centrifuged at 800*g* for 10 min at 4 °C. Supernatants were collected, transferred to fresh 1.5 mL tubes, and stored at −80 °C. Total spleen, total large intestine, total lungs and 200 to 250 mg of small intestine were suspended in 300 µL of PBS in pre-filled 2.0 mL tubes containing zirconium beads (cat 155-40945, Spectrum). Tubes were processed in a BeadBug-6 homogenizer at a speed setting of 3,000 and a 30 second cycle time for four cycles with a 30-second break after each cycle. Tissue homogenates were centrifuged at 15,000 rpm for 20 minutes at 4 °C. Homogenate supernatants were then transferred into 1.5 mL tubes and stored at −80 °C. STI-9167 antibody levels in each sample were quantified using the antibody detection ELISA method. Statistical significance was determined using the Welch’s t-test. This study was reviewed and accepted by the animal study review committee (SRC) and conducted in accordance with IACUC guidelines.

### Pharmacokinetic Study

Female CD-1-IGS (strain code #022) were obtained from Charles River Laboratories at 6-8 weeks of age. STI-9167 was dissolved in intranasal formulation buffer C was administered as previously described for the IN biodistribution study. Lungs and blood were collected from 6 mice at each of the following timepoints: 10 min, 1.5 h, 6 h, 24 h, 72 h, 96 h, 168 h, 240 h, and 336 h. Serum and lung tissue samples were collected as described for the biodistribution study. STI-9167 antibody levels in each sample were quantified using the antibody detection ELISA method. Pharmacokinetic analysis of the collected ELISA data was performed with the Phoenix WiNnonlin suite of software (version 6.4, Certara) using a non-compartmental approach consistent with an IN-bolus route of administration. Statistical significance was determined using the Welch’s t-test. This study was reviewed and accepted by the animal study review committee (SRC) and conducted in accordance with IACUC guidelines.

### khACE2 mouse model of COVID-19 infection

K18-hACE2 transgenic mice were purchased from Jackson laboratory and maintained in pathogen-free conditions and handling conforms to the requirements of the National Institutes of Health and the Scripps Research Institute Animal Research Committee. 8-12 weeks old mice were infected intranasally with 10,000 PFU of SARS-COV-2 in total volume 50 μL different concentration of AB were injected intravenously 1 h post infection or by intranasal instillation 12 h post infection.

### Determination of infectious virus titer in the lung

On day 4 post-infection, animals were euthanized, lung tissue samples were collected from each animal, and the left lobe of each collected lung was placed into a pre-labeled microcentrifuge tube containing 3-5 beads 2.3 mm diameter Zirconia/silica beads (Fischer). Lung samples were homogenized with DMEM + 5% FBS in a TissueLyser 1 min 25 sec ^55^.

VeroE6 cells were plated at 3.0E+055 cells/well in 24 well plates in volume 400 μL/well. After 24 h. medium was removed, and serial dilution of homogenized lungs were added to Vero cells and subsequently incubated for 1 h at 37 °C. After incubation, an overlay (1:1 of 2% methylcellulose [Sigma] and culture media) is added to each well and incubation commenced for 3 days at 37 °C. Plaque staining was performed using Crystal Violet as mentioned above. Virus titers in lungs were compared with the isotype control mAb-treated group using a Student’s t-test.

### Plaque reduction neutralizing assay

VeroE6 cells were plated at 18.0E+03 cells/well in a flat bottom 96-well plate in a volume of 200 μL/well. After 24 h, a serial dilution of ABs is prepared in a 100 μL/well at twice the final concentration desired and live virus was added at 1,000 PFU/100 μL of SARS-COV-2 and subsequently incubated for 1 h at 37 °C in a total volume of 200 μL/well. Cell culture media was removed from cells and sera/virus premix was added to VeroE6 cells at 100 μL/well and incubated for 1 h at 37 °C. After incubation, 100 μL of “overlay” (1:1 of 2 % methylcellulose (Sigma) and culture media) is added to each well and incubation commenced for 3 d at 37 °C. Plaque staining using Crystal Violet (Sigma) was performed upon 30 min of fixing the cells with 4% paraformaldehyde (Sigma) diluted in PBS. Plaques were assessed using a light microscope (Keyence).

## ACKNOWLEDGEMENTS

This work was partly supported by CRIPT (Center for Research on Influenza Pathogenesis and Transmission), an NIAID funded Center of Excellence for Influenza research and Response (CEIRR, contract #75N93021R00014 to AGS, MS, FK), by DARPA grant HR0011-19-2-0020 (to AGS) and by NCI Seronet grant U54CA260560 (to AGS, MS, FK). We thank R. Albrecht for support with the BSL-3 facility and procedures at the Icahn School of Medicine at Mount Sinai, NY. S.Y. received funding from Swiss National Foundation (SNF) Postdoc Mobility fellowship (P400PB_199292). M.S. laboratory is supported by NIH grant R01DK130425.

## FIGURE LEGENDS

**Supplemental Figure 1.**
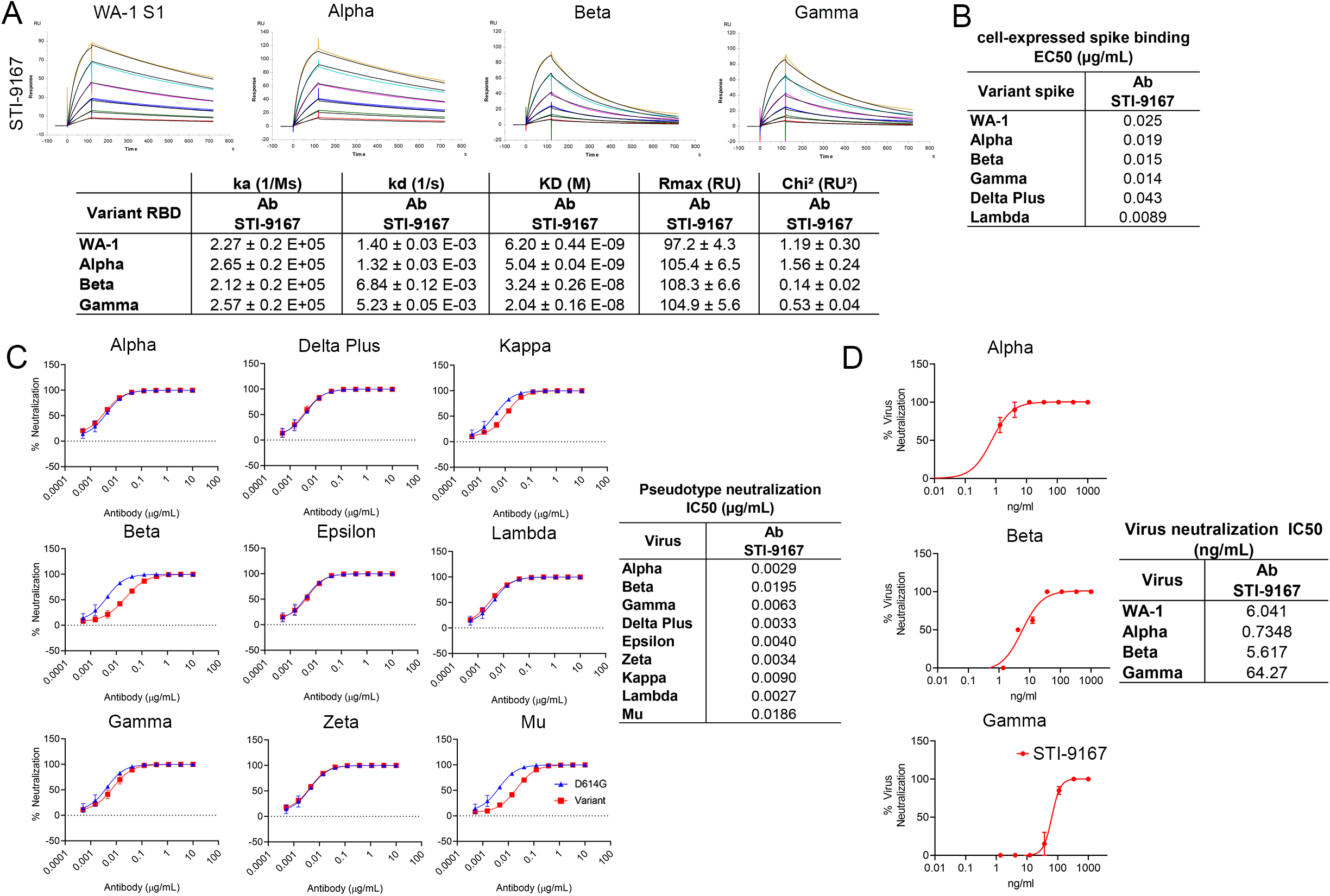
Binding and neutralization of candidate antibody to VoCs. **(A)** Affinity measurements of STI-9167 for Spike S1 binding domain from the following isolates and VOCs: USA/WA-1/2020(WA-1) isolate, Alpha, Beta, and Gamma. The antibody affinities were measured using SPR on a BIAcore T200 instrument using a 1:1 binding model. Graphs are representative of triplicate data and table data presented as mean ± SD. **(B)** Spike protein derived from Alpha, Beta, Gamma, Delta Plus, and Lambda SARS-CoV-2 isolates were independently expressed on the surface of HEK 293 cells. Serially-diluted STI-9167 was assayed for Spike protein binding by flow cytometry. To quantify antibody binding, mean fluorescent intensity was measured for each dilution tested and the EC_50_ value was calculated for each nAb. **(C)** Spike-pseudotyped VSV neutralization. Antibody neutralization of the indicated spike variant pseudotyped VSVs was performed as described in the methods. The curves represent the average of three independent experiments, with error bars representing one standard deviation. IC50 values for each pseudotype/antibody combination are indicated on the right. **(D)** PRNT assay using STI-9167 with indicated SARS-COV-2 variants were performed as described in the methods.

**Supplemental Figure 2.**
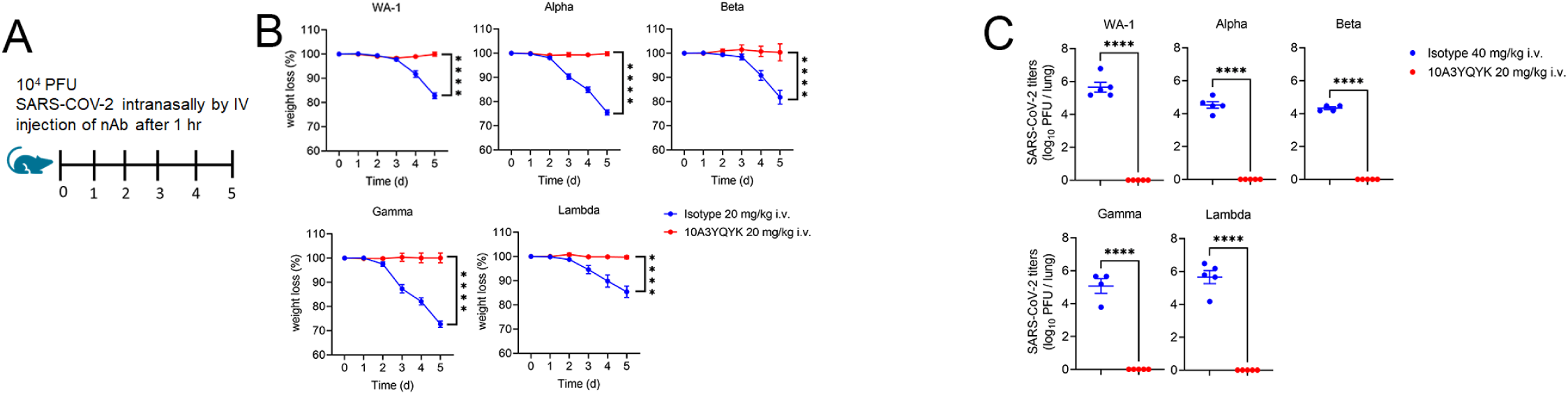
Efficacy of Intranasal (IN) delivery of STI-9167 Neutralizing Antibody in the K18-hACE2 murine model of COVID-19 VoCs. **(A)** A schematic of experimental model, K18-hACE2 transgenic mice were infected with 10000 PFU of indicated variants of SARS-CoV-2 treated with indicated concentration of AB (STI-9167) intravenously 1 hour post infection. **(B)** Body weight change of mice was measured daily (n = 5). **(C)** SARS-CoV-2 viral titers were measured in lung day 5 post infection (n = 5). P****<0.0001. Unpaired t test (C). Two-way ANOVA (B)

**Supplemental Figure 3.**
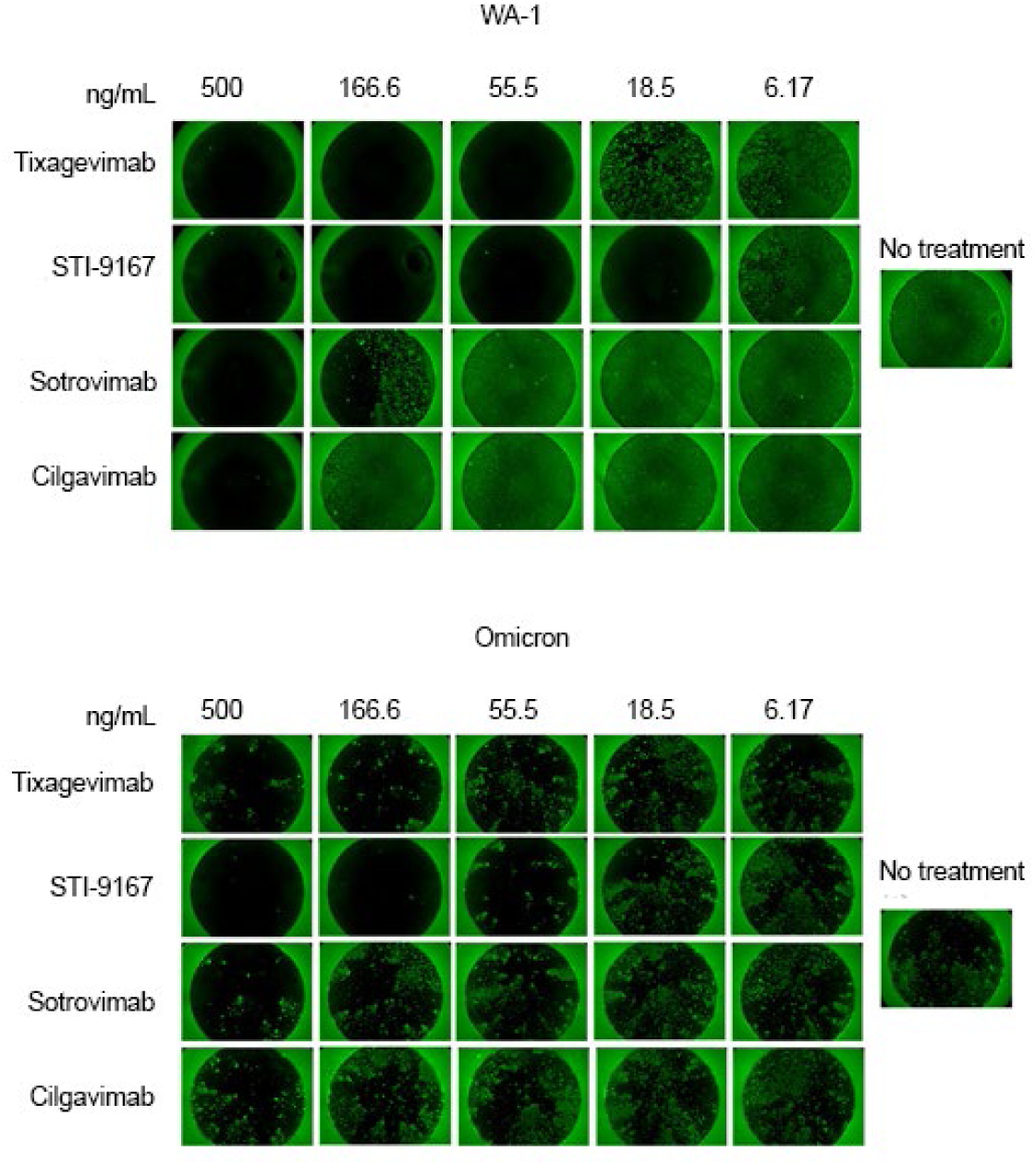
Plaque Reduction Neutralization Fluorescent staining on Vero-ACE2 cells. PRNT assay using STI-9167 and various neutralizing antibodies with SARS-COV-2 WA-1 or Omicron were performed as described in the methods on Vero-ACE2-expressing cells and visualized.

**Supplemental Figure 4.**
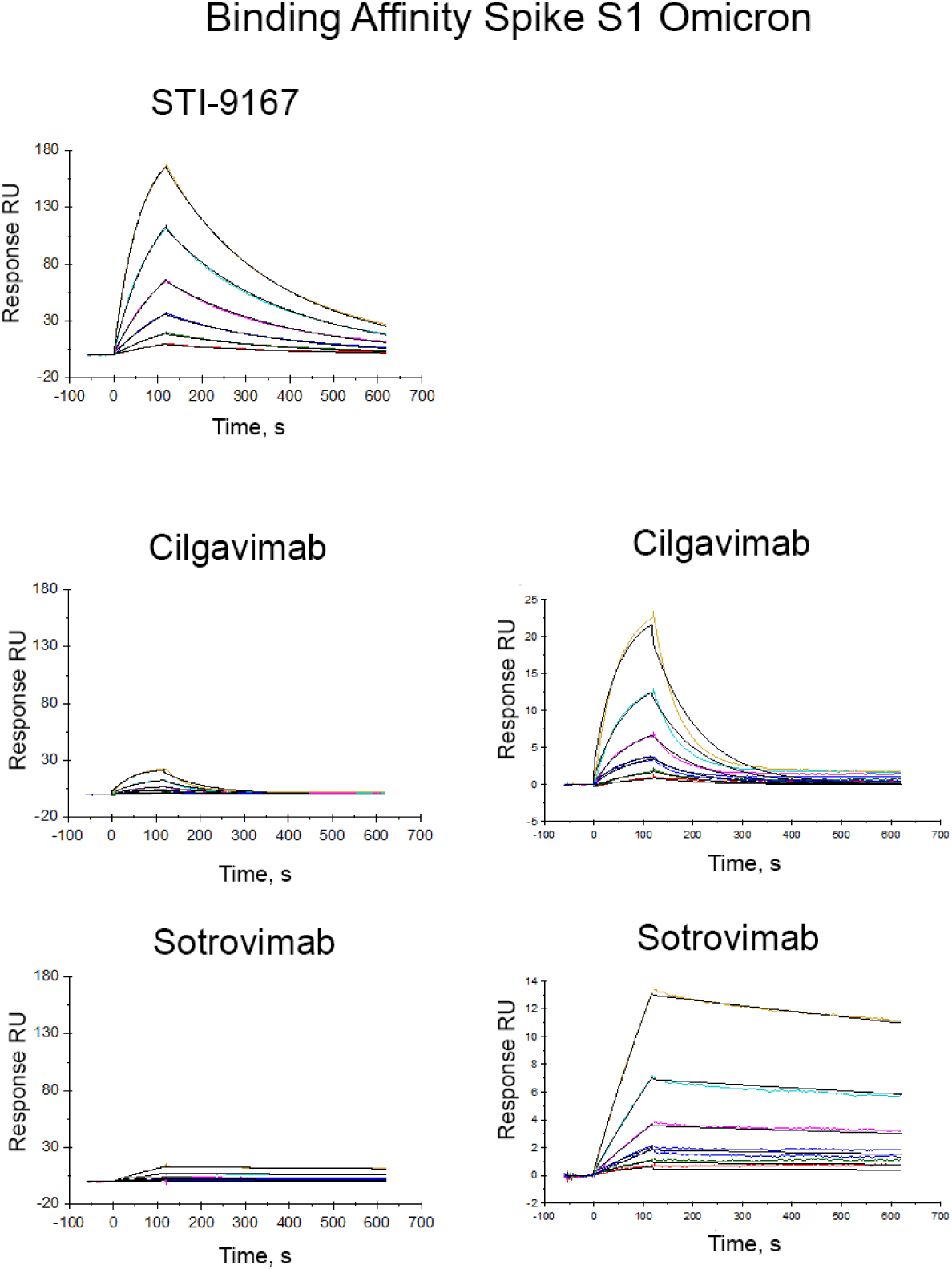
Binding Affinity of Neutralizing Antibodies to Omicron Spike Protein. SPR binding affinity graphs of STI-9167, Cilgavimab, Tixagevimab, and Sotrovimab.

**Supplemental Figure 5.**
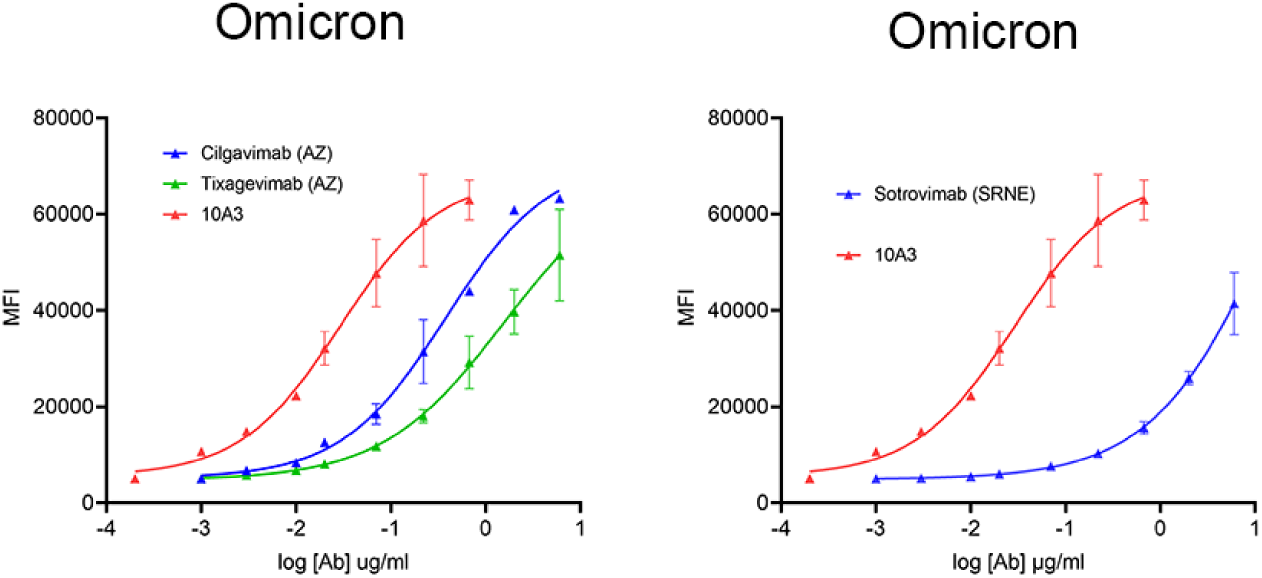
Cell-expressed spike binding to Neutralizing Antibodies. Omicron spike protein was expressed on HEK 293 cells and binding of selected neutralizing antibodies was measured by MFI.

